# A global regulatory atlas of *Streptomyces* reveals conserved and diversified transcriptional networks across actinomycetes

**DOI:** 10.64898/2026.06.06.730586

**Authors:** Hannah E. Augustijn, Hiroshi Otani, Augustin Rigolet, Rachel Stuij, Vina My Pham, Leo Baumgart, Le Zhang, Yu Zhang, Chao Du, Simona Cernat, Sébastien Rigali, Ronan O’Malley, Marnix H. Medema, Nigel J. Mouncey, Gilles P. van Wezel

**Author notes:** These authors contributed equally (Hiroshi Otani, Augustin Rigolet). Department of Chemistry and Chemical Biology, Harvard University, Cambridge, MA, USA. The University of Chicago, Division of the Biological Sciences, Chicago, IL, USA.

## Abstract

Transcriptional regulatory networks determine how bacteria integrate environmental signals with growth, metabolism and stress adaptation. Actinomycetes encode exceptionally large repertoires of transcription factors (TFs) coordinating morphological development, environmental adaptation and specialized metabolism, yet their regulatory networks remain poorly defined. Here, we apply DNA affinity purification sequencing (DAP-seq) to 789 predicted TFs of *Streptomyces coelicolor*, generating genome-wide binding maps for 393 regulators and expanding the experimentally supported regulome from ∼8% to ∼50%. Integration with ChIP-seq reveals pleiotropic regulators and hierarchical network architecture linking primary metabolism, development and biosynthetic gene clusters (BGCs). Multiplexed DAP-seq (multiDAP) across 16 additional actinomycetes uncovered deeply conserved regulatory circuits alongside widespread divergence in TF-target interactions. Together, these data establish a regulatory atlas for *Streptomyces* and related actinomycetes, enabling researchers to explore control of genes, pathways, BGCs and conserved network modules. This provides a foundation for predictive analysis and engineering of complex bacterial phenotypes in biotechnology and medicine.

## INTRODUCTION

Twenty-five years ago, the complete genome sequence of *Streptomyces coelicolor* transformed our perception of actinomycete genetics ^1^. Until then, the biosynthetic and developmental capacity of these bacteria was inferred largely from observable phenotypes. Genome sequencing revealed the scale of regulatory and metabolic complexity encoded in actinomycete genomes, including dozens of previously unrecognized biosynthetic gene clusters (BGCs), many of them silent under laboratory conditions. This discovery uncovered a vast reservoir of hidden chemical potential and reshaped natural product research by showing that metabolic capacity far exceeds phenotypic output ^2,3^. Today, a comparable blind spot exists at another layer of biological organization: transcription factor regulatory networks (TFRNs) that govern development, stress adaptation and specialized metabolism in actinomycetes ^4^.

Actinomycetes possess some of the most complex and intertwined regulatory systems in bacteria. The genome of the model organism *S. coelicolor* encodes ∼900 transcription factors (TFs) and sigma factors, representing ∼12% of the genome ^1^. This unusually large regulatory repertoire reflects the need to coordinate multicellular development, environmental sensing and secondary metabolism ^3^. Yet binding sites have been experimentally identified for only a small fraction of these regulators, leaving a major gap in our understanding of regulatory architecture ^5,6^. In contrast, model organisms such as *Escherichia coli, Pseudomonas aeruginosa* and *Bacillus subtilis* benefit from near-complete regulatory maps that enable systems-level modeling and predictive manipulation of gene expression ^7–9^. The absence of such a regulatory atlas in actinomycetes limits interpretation of genome content, constrains activation of silent BGCs and hampers evolutionary analysis of regulatory architecture.

Streptomycetes are multicellular mycelial bacteria that undergo a complex developmental program culminating in sporulation ^10,11^. Streptomycetes produce a wide range of bioactive natural products, including antibiotics, antifungals, anticancer agents and immunosuppressants ^12,13^ and encode extensive enzymatic repertoires that support their saprophytic lifestyle ^14^. Historically, most well-characterized *Streptomyces* regulators were discovered through phenotype-driven genetics, including Whi (white) mutants that fail to produce grey spore pigment ^15^, Bld (bald) mutants that fail to form aerial hyphae ^16^, and mutants that lost the ability to synthesize pigmented antibiotics (Red) ^17^. While this classical approach yielded foundational insights, it is inherently biased toward regulators that produce visible phenotypes under laboratory conditions.

Regulatory network information provides a powerful and largely untapped dimension for genome interpretation. In specialized metabolism, integration of TF binding data with gene cluster analysis can predict BGC function, which enabled discovery of a previously unrecognized locus required for desferrioxamine biosynthesis ^18^. These examples suggest that TFRNs represent a systems-level layer of information whose impact is analogous to that of genome sequencing. However, systematic reconstruction of these networks has been constrained by the lack of scalable methods to interrogate transcription factors at genome scale. Chromatin immunoprecipitation sequencing (ChIP-seq) enables genome-wide mapping of individual TFs *in vivo* ^19,20^. DNA affinity purification sequencing (DAP-seq) provides a scalable *in vitro* alternative, interrogating genome-wide binding by all TFs encoded by the genome ^21,22^, while multiplexed DAP-seq (multiDAP) ^21^ enables taxonomic level determination of conservation and uniqueness of regulatory functions. A recent study applied DAP-seq specifically to study binding of putative gamma-butyrolactone-binding cluster-situated regulators (CSRs) across streptomycetes, showing autoregulation of BGCs ^23^. To move from regulator-by-regulator discovery to systems-level understanding of transcriptional control in *Streptomyces*, and obtain insights into the global regulatory architecture encoded by the full transcription factor repertoire, a genome-scale reconstruction of transcriptional regulatory networks is required.

Here, we establish a scalable strategy for global reconstruction of transcriptional regulatory networks in *Streptomyces*. Using DAP-seq, we profiled 789 predicted transcription factors of *S. coelicolor* and obtained high-confidence genome-wide binding profiles for 393 regulators, hugely expanding the experimentally supported regulome. This unbiased, phenotype-independent approach reveals extensive pleiotropy and a hierarchical network architecture linking primary metabolism, development, stress responses and biosynthetic gene clusters. To enable cross-species comparison, we applied multiDAP under standardized conditions across 16 additional actinomycetes, revealing deep conservation of core regulatory circuits alongside lineage-specific changes in TF–target connectivity. Together, these data establish a regulatory atlas for comparative and predictive analysis of transcriptional control, providing a foundation for understanding and engineering complex bacterial phenotypes in actinomycetes.

## RESULTS

### Genome-scale reconstruction of transcriptional regulatory architecture in Streptomyces coelicolor

To enable genome-scale reconstruction of transcriptional regulatory networks, we aimed to characterize the binding of all transcription factors (TFs) of the model organism *S. coelicolor* M145, generating constructs for 789 of 864 target regulators to perform DAP-seq (Fig. 1a, Table S1). This allows the identification of TF target genes *in vitro*. All peak data are freely available through the StrepTRN data portal, accessible at: https://streptrn.bioinformatics.nl.

**Fig. 1:**
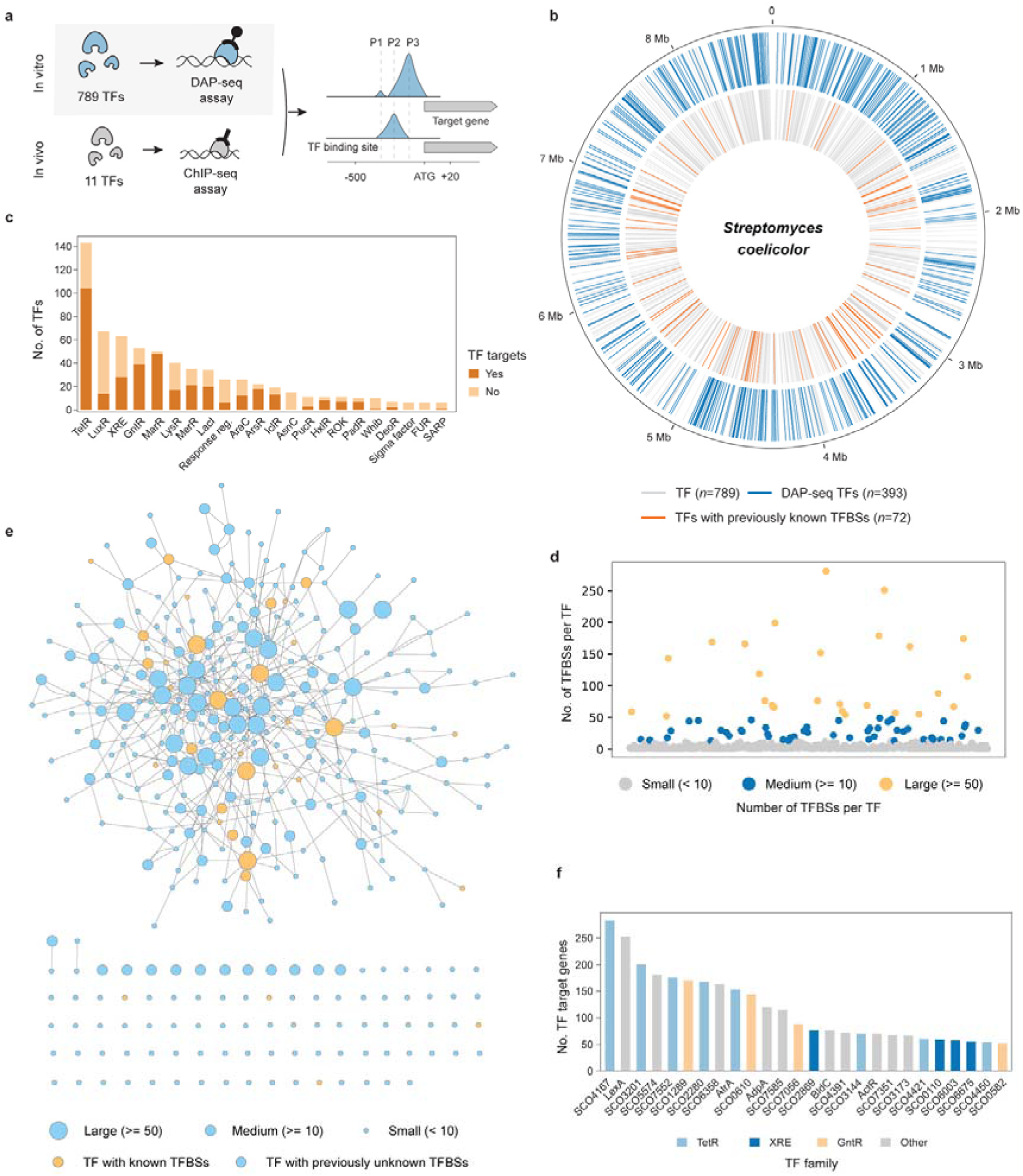
Expansion of the regulatory network of *S. coelicolor*. **a,** Schematic overview of the 789 TFs tested by DAP-seq and 11 TFs additionally analyzed by ChIP-seq. Peaks (P) were assigned to target genes using a window from -200 to +20 relative to the start codon. **b,** Overview of TF genes on the *S. coelicolor* chromosome. Grey bars indicate genes encoding putative TFs with no identified binding sites; blue outer bars show genes for TFs with newly identified DAP-seq TFBSs; orange inner bars denote TF genes associated with previously characterized TFBSs. **c,** Number of TFs per family, showing only families with at least 10 members that have demonstrated binding sites passing the filtering criteria. **d,** Scatter plot showing the distribution of TFBSs per TF. The fractions of TFs are shown for small (<10, grey), medium (10–50, blue), and large (>50, yellow) numbers of TFBSs. **e,** Regulatory network of TF cross-regulation. Nodes represent TFs; edges indicate TFBSs identified upstream of other TFs. Node size reflects the number of TFBSs (small, medium, large), and orange nodes denote TFs with previously characterized binding sites. **f,** Number of target genes for TFs with large TFBSs numbers.

We identified high-confidence binding sites for 393 TFs using a minimum fraction of reads in peaks score of 0.05, positions spanning −200 to +20 relative to the start codon, and a ≥30-fold enrichment over background (see Online Methods). This represents a substantial increase compared to the ∼72 TFs for which at least one binding site had been previously validated experimentally (Fig. 1b, Table S2). This expands the experimentally supported regulome from approximately ∼8% to ∼50% of all *S. coelicolor* TFs. Moreover, whereas prior studies have typically focused on single gene targets, our analysis assessed genome-wide TF binding sites (TFBSs), and identified 5,425 TFBSs, representing an estimated tenfold increase in experimentally identified sites.

The 393 DAP-seq TFs belong to 23 major TF families, among which the MarR, ArsR, and TetR families showed notably high recovery rates of 98%, 83%, and 71%, respectively (Fig. 1c; Table S1). The absence of detectable binding for certain TFs or TF families is consistent with known *in vitro* limitations, as they may require activation mechanisms or ligand binding that are not present under standard DAP-seq conditions ^21,22^. Additionally, TF binding in DAP-seq may be limited by technical factors such as protein misfolding or instability, and reduced detection of low-affinity or transient interactions.

Analysis of TFBS distributions revealed that most TFs (77%, *n* = 302) have ten or fewer TFBSs, with 33% (*n* = 128) having only a single detected binding site (Fig. 1d). In contrast, 7% (*n* = 26) had 50 or more TFBSs, suggesting a potential role as global regulators. Mapping TF cross-regulatory interactions revealed a highly connected network centered around TFs with large regulons that bind upstream of other TFs, indicating multi-layered regulatory control, many of which remain uncharacterized (Fig. 1e). Together, these features reveal a hierarchical and highly interconnected network, rather than isolated regulons. The largest regulons were observed for SCO4167, a TetR-family regulator with 281 TFBSs, and the DNA damage response regulator LexA (SCO5803), with 251 binding sites (Fig. 1f). While LexA has been extensively studied in *Streptomyces* ^24,25^, the function of SCO4167, along with many other TFs, remains largely uncharacterized, highlighting the potential of these data to enable systematic exploration of previously unknown regulators.

DAP-seq also serves as an unbiased screen to capture both canonical and non-canonical DNA binding. For example, while most response regulators (RRs) are phosphorylation-dependent, binding was detected for orphan TFs lacking their cognate histidine kinase, including RamR ^26^, GlnR ^27^, and the uncharacterized SCO3144 (69 TFBSs). Notably, several DAP-seq TFBSs overlap with sites previously reported in the RamR and GlnR studies. TFBSs were also identified for the non-orphan response regulators SCO7230 and SCO2143, suggesting potential binding in an unphosphorylated state.

### Validation of the DAP-seq data via ChIP-seq

To validate the DAP-seq data, we compared our results to previously characterized TFs. For eight TFs with sufficient data (at least four sites in both datasets), sequence motifs showed high concordance with published motifs, confirming that DAP-seq captures established binding specificities (Fig. 2a). We further assessed concordance between *in vitro* and *in vivo* binding by performing ChIP-seq for 11 representative TFs (Fig. 1a, 2b, Table S3). DAP-seq captured a substantial subset for most TFs (Fig. 2b). A notable exception was the ArsR-family regulator SCO3696; no binding was detected *in vitro*, whereas ChIP-seq identified clear *in vivo* targets, indicating a requirement for physiological context or cofactors. Conversely, DAP-seq identified additional candidate targets for some TFs, such as the heat shock regulator HspR (SCO3668) ^28^.

**Fig. 2:**
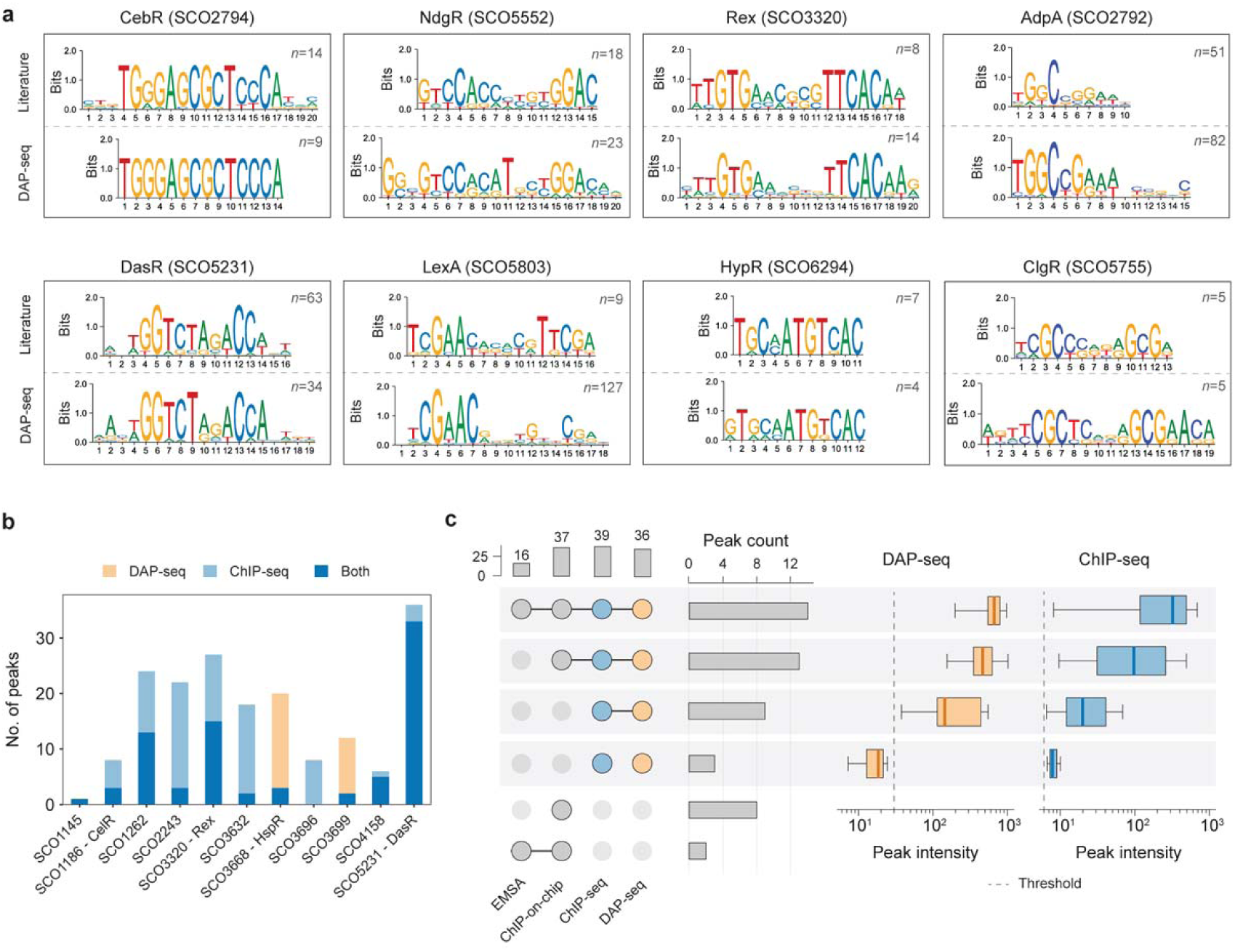
Validation of the DAP-seq. **a,** Comparison of DAP-seq versus literature derived sequence motifs. **b,** Comparison of the number of peaks retained in DAP-seq and ChIP-seq analyses. **c,** Comparison of DasR (SCO5231) regulons identified by DAP-seq and ChIP-seq with previously published ChIP-on-chip and EMSA data ^30^. Box plots show peak intensities relative to a background distribution. The dotted line indicates a ≥30-fold enrichment cutoff for DAP-seq, separating low-confidence peaks from high-confidence peaks.

Comparison of DAP-seq and ChIP-seq profiles for the well-characterized DasR (SCO5231) ^29,30^ showed that 27 out of 37 (73%) of the previously identified targets were recovered by both approaches (Fig. 2c, Table S4). The large proportion of high-intensity DAP-seq peaks among ChIP-supported targets indicates that stronger DAP-seq signals reflect high-confidence binding, consistent with previous cross-platform validation studies ^21^.

### Functional assignment of transcription factors

The majority of identified TF–target interactions remain functionally uncharacterized, limiting the interpretation of their biological role and potential exploitation. To infer TF function, we linked target genes to KEGG and COG annotations and performed an enrichment analysis to identify regulators associated with specific metabolic and cellular pathways ^31,32^. This revealed 20 TFs significantly associated (Benjamini-Hochberg false discovery rate [FDR] *P*_adj_□<□0.05) with specific pathways or functional categories, including amino acid, carbohydrate, energy & carbon and lipid metabolism (Fig. 3a). Of these TFs, 12 have previously predicted regulons or high sequence similarity to characterized orthologs. The TF most strongly associated with the cell cycle pathway (KEGG) is SCO1383 (Fig. 3a,b), which shares low amino acid sequence identity (43%) with FtsR from *Corynebacterium glutamicum* ^33^. In *C. glutamicum*, FtsR regulates cell division by binding upstream of *ftsZ*, a critical component of Z-ring formation and cytokinesis ^34,35^. As discussed in the multiDAP section below, SCO1383 is extremely well conserved across actinomycetes. Similarly, the TF with the second strongest functional enrichment, SCO1171, is associated with pentose and glucuronate interconversions (Fig. 3a,c). Indeed, SCO1171 is the predicted ortholog of XylR, a repressor of D-xylose metabolism characterized in *Streptomyces avermitilis* ^36^. DAP-seq data analysis revealed SCO1171 binding within the divergent *xylAB* promoter region (SCO1169-1170), and transcriptomic analysis confirmed induced expression of the SCO1169-1171 cluster under the xylose condition compared to the glucose condition (FDR *P*_adj_ < 0.01; Fig. 3d) ^37^.

**Fig. 3:**
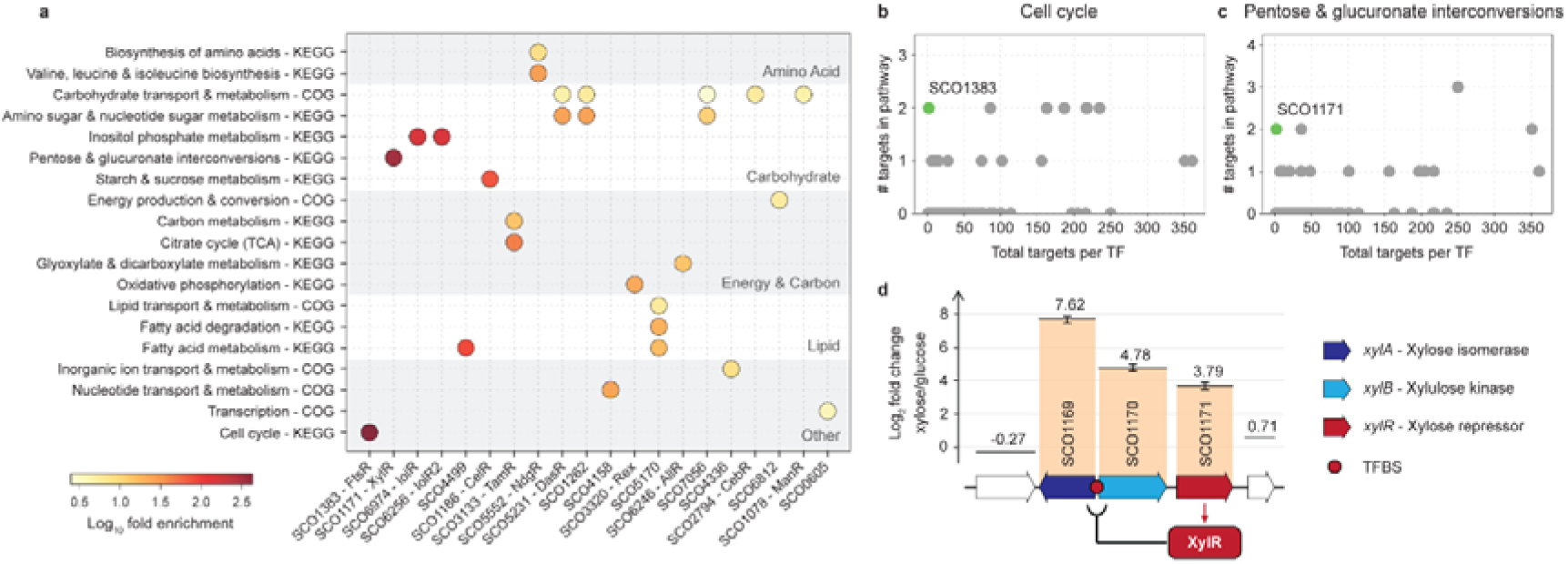
Functional assignment of *S. coelicolor* TFs. **a,** Gene set enrichment for KEGG metabolic pathways and COG functional categories. Enrichment was assessed via the Fisher’s exact test with Benjamini-Hochberg correction (FDR *P*_adj_□<□0.05); color intensity indicates relative enrichment. **b,c** Gene enrichment plot showing total target genes per regulator versus targets in cell cycle pathways **(b)** or pentose & glucuronate interconversions **(c)**. Non-significant results (FDR *P*_adj_ ≥□0.05) are in grey. **d,** Regulon for XylR (SCO1171). Significantly differentially expressed genes (FDR *P*_adj_ < 0.01 or |log₂FC| > 1) of *S. coelicolor* grown in xylose vs glucose as carbon source ^37^ are indicated by colored bars with error bars representing mean ± s.d. (*n* = 3). Statistical analysis was performed using an unpaired Welch’s t-test with Benjamini-Krieger-Yekutieli FDR correction. Values represent mean expression changes.

Additionally, we identified eight previously uncharacterized TFs associated with distinct metabolic and cellular processes, namely fatty acid metabolism (SCO4499, SCO5170), carbohydrate metabolism (SCO1262, SCO7056), nucleotide transport (SCO4158), inorganic ion transport (SCO4336), energy metabolism (SCO6812), and general transcription (SCO0605). Together, these findings recover known regulatory relationships and uncover previously unrecognized regulators, providing a framework for targeted exploration of transcriptional control.

### Regulatory dynamics of BGC expression

Unlocking the metabolic potential of actinomycetes requires a systematic understanding of the regulatory networks governing silent BGCs. We mapped TF binding across antiSMASH-predicted BGCs in *S. coelicolor* ^38^, using refined boundaries based on MIBiG references ^39^, co-expression analysis ^18^, or manual annotation of core biosynthetic genes (Table S5, S6). This identified 51 TFs targeting promoter regions of 16 out of 27 clusters, twelve of which represent or share high similarity (≥90%) to BGCs encoding the production of characterized compounds (Fig. 4a).

**Fig. 4:**
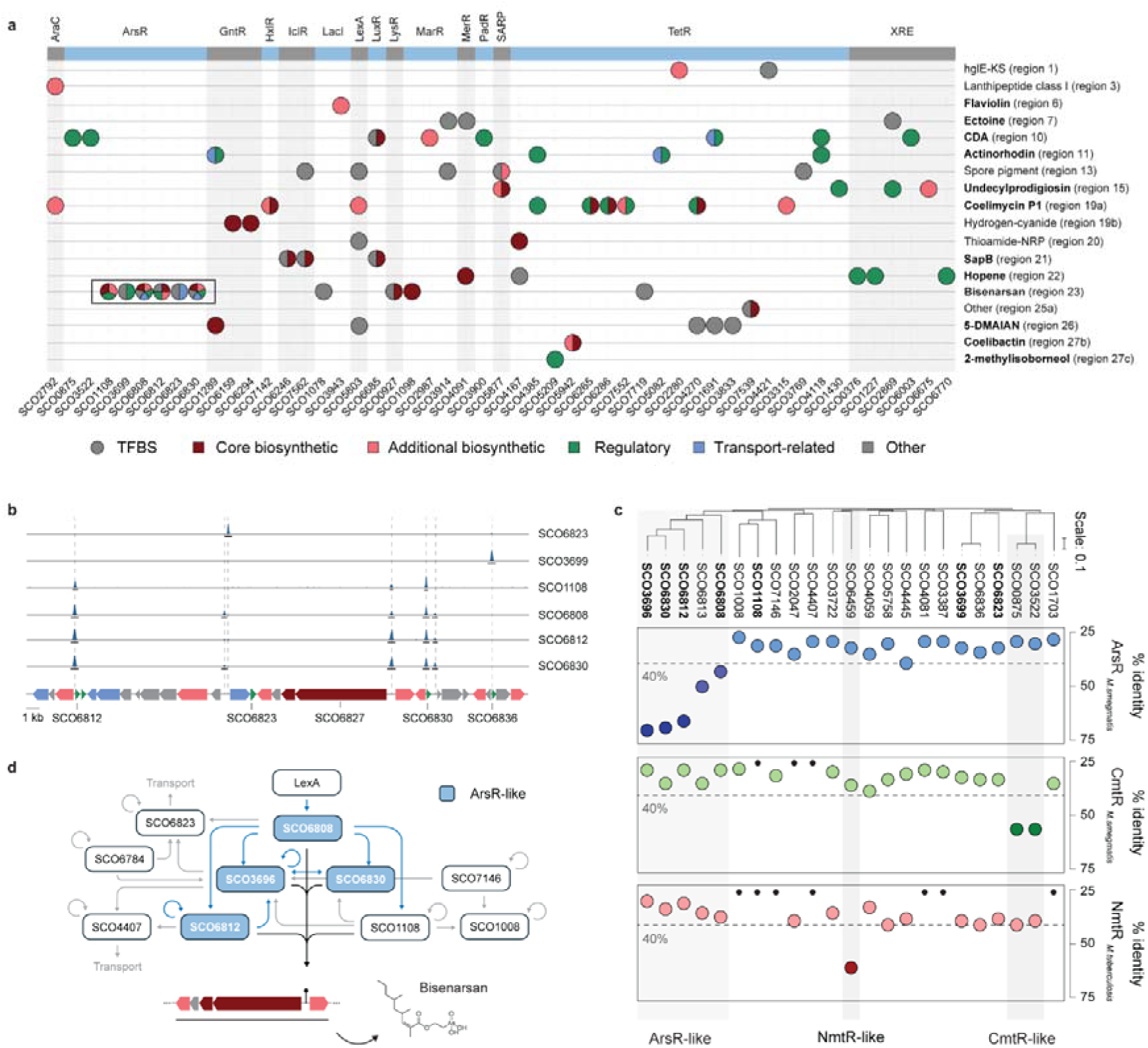
Regulation of *S. coelicolor* BGCs. **a,** Overview of TF binding within promoter regions of antiSMASH-predicted BGCs. Each dot represents a TF binding site, with TFs grouped by family. BGC identifiers correspond to antiSMASH v7 region numbers; colored circle segments indicate functional annotations of genes adjacent to predicted TF binding sites, based on gene orientation. A complete list of TF binding events is provided in Table S3. **b,** DAP-seq binding of ArsR-family regulators within BGC region 23. **c,** Sequence similarity of *S. coelicolor* ArsR-family TFs to ArsR and CmtR from *M. smegmatis* and NmtR from *M. tuberculosis*. **d,** Regulatory network of the bisenarsan BGC as inferred from DAP-seq data.

To resolve regulatory logic at the cluster level, we examined BGC region 23, the *S. coelicolor* homolog of the arsenate-induced bisenarsan cluster ^40^. DAP-seq revealed six ArsR-family regulators binding within this cluster (Fig. 4b), a family typically defined by metal-responsive ligands such as ArsR (arsenite/antimonite), CmtR (cadmium/lead), SmtB (zinc), and KmtR/NmtR (nickel/cobalt). Five regulators (SCO3696, SCO6830, SCO6812, SCO6808, and SCO6813) share significant homology with the *Mycobacterium smegmatis* ArsR ortholog ^41^ (Fig. 4c). Although SCO3696 binding was not detected *in vitro*, ChIP-seq confirmed overlapping *in vivo* targets of SCO3696 binding upstream of SCO6830 and SCO6812 (Table S3), and motif analysis supports conserved ArsR-like binding (Fig. S1). The regulatory profile of these ArsR-like regulators matches the known arsenate-induced production of bisenarsan ^40^, indicating that they are likely arsenite/antimonite responsive. By contrast, SCO1108 binds directly upstream of the bisenarsan BGC, but its physiological role remains unclear. Thus, the bisenarsan cluster exemplifies BGC control by hierarchical, interconnected regulatory networks rather than single regulatory switches (Fig. 4d). This network extends beyond biosynthesis to transport, resistance and stress-response control. ArsR-family regulators target transporter-related genes, while LexA binding upstream of SCO6808 links bisenarsan biosynthesis directly to the SOS response. Thus, BGC expression is embedded within global stress-response networks rather than controlled by isolated, cluster-specific regulators. This logic extends to other BGCs: the pleiotropic regulator AdpA/BldH (SCO2792) ^42^ binds upstream of SCO0268, encoding a putative ribosomally synthesized and post-translationally modified peptide (RiPP) precursor within an uncharacterized class I lanthipeptide cluster (BGC region 3). Together, these findings show that genome-wide TF binding maps reposition BGC regulation within multilayered regulatory hierarchies and provide a rational route for targeted activation of silent biosynthetic pathways.

### MultiDAP highlights regulatory network conservation across actinomycetes

To investigate how transcriptional regulatory networks are conserved and changed, and to validate predicted TF-target interactions through recurrent binding at orthologous loci, we compared TF-target interactions across 17 taxonomically diverse actinomycetes (Table S7). Using multiDAP, the 789 affinity-purified *S. coelicolor* TFs were incubated with barcoded genomic libraries, generating a dataset equivalent to >13,000 individual DAP-seq experiments and identifying 110,539 TF binding sites across species under identical *in vitro* conditions. Binding profiles for various regulators aligned with previously characterized binding profiles, including the copper-sensing CsoR and the p-hydroxybenzoate hydroxylase regulator PobR ^43,44^, validating the cross-species accuracy of our identified sites (Fig. 5a).

**Fig. 5.**
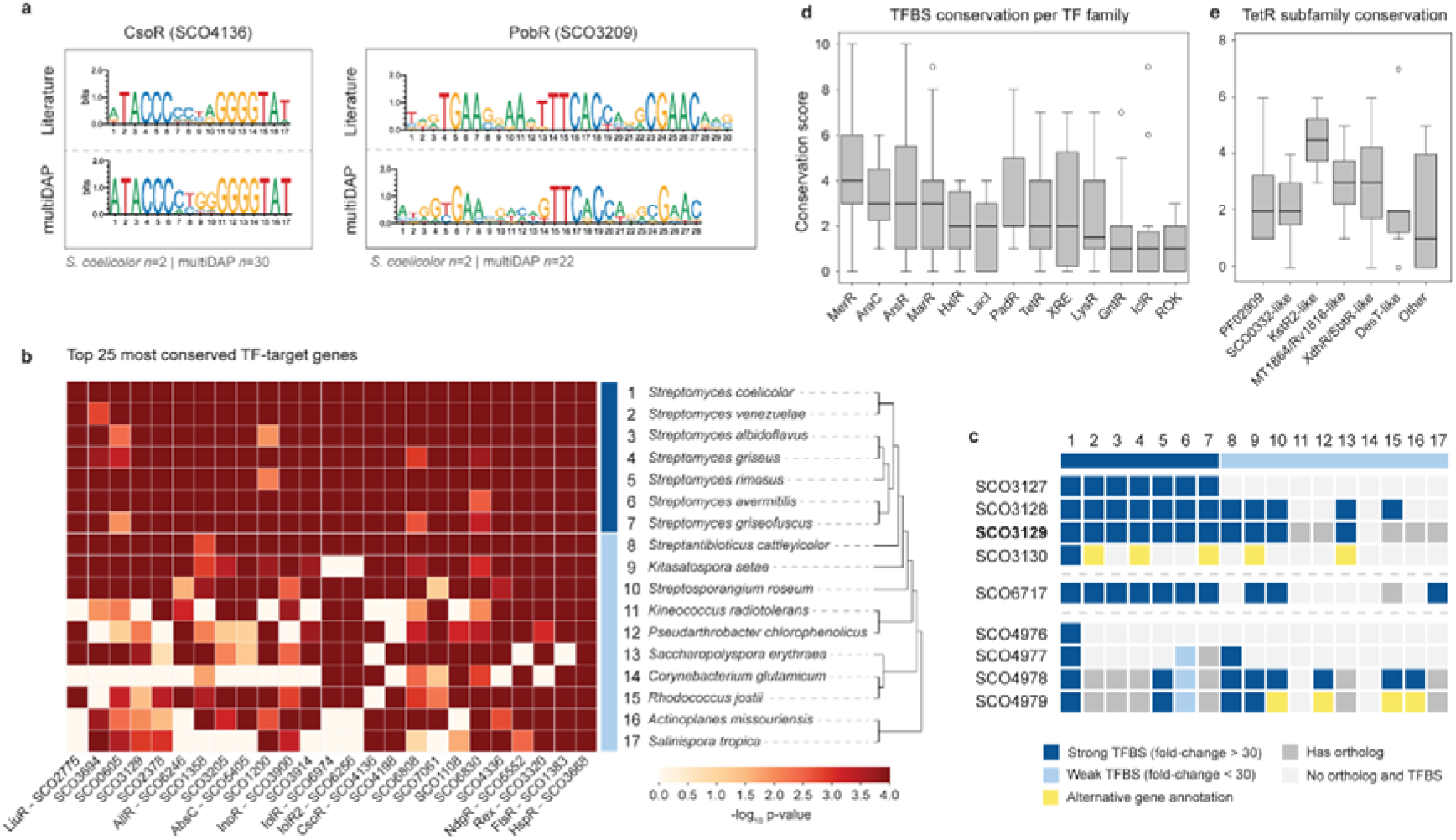
MultiDAP reveals conserved and rewired regulatory interactions across 17 actinomycetes. **a,** comparison of consensus sequences for the binding sites of CsoR and PobR between literature and multiDAP data. **b,** The 25 most conserved TF–target interactions identified by multiDAP across 17 actinomycetes. Conservation was defined by binding of *S. coelicolor* TFs to orthologous loci in the other 16 genomes. Species are grouped by phylogeny, with members of the *Streptomyces* clade shown in dark blue (species 1–7) and non-*Streptomyces* spp. shown in light blue (species 8–17). The color scale reflects statistical enrichment, with the strongest conservation indicated by *P*□≤□1□×□10⁻^4^. **c,** Conservation of SCO3129 target genes across 17 actinomycetes. Strong (dark blue) and weak (light blue) TFBSs are shown per species; grey indicates orthologs without binding, white absence of both ortholog and TFBS, and yellow alternative gene annotations. Conserved binding at SCO3128, SCO4978 and SCO6717 indicates a shared role in fatty acid metabolism and membrane homeostasis, whereas variation at other loci suggests regulatory rewiring. **d,** Degree of conservation (*P* < 0.01) of *S. coelicolor* TFBSs across the 10 non-*Streptomyces* species, grouped by TF family. **e,** TetR subfamily determined based on protein domain architecture. Each dot represents a single conserved TF target.

To identify conserved regulatory networks, we mapped TF binding sites to orthologous genes across the 17 genomes and validated the frequency of these sites against a null model that accounts for lineage-specific variations in regulon size ^21^. This identified a core of highly conserved TF-operon interactions (Fig. 5b). The five most highly conserved binding profiles were associated with the heat-shock regulator HspR that binds at the *dnaK* and *clpB* chaperone promoter regions ^28,45,46^, the cell division regulator SCO1383 that binds to the promoter region of the cell division scaffold gene *ftsZ*, the redox sensor Rex ^47^, the amino acid regulator NdgR ^48^ and SCO4336. SCO1383 bound immediately upstream of *ftsZ* in 16 of the 17 actinomycetes, supporting its role as a *bona fide* regulator of *ftsZ* in streptomycetes. Indeed, we confirmed the binding via EMSA experiments (Fig. S2). SCO4336 is a MarR-family regulator whose predicted targets include genes for the tRNA modifying enzymes SCO2497 and SCO5164, for an Na+/H+ antiporter (SCO3185), and several *marR* regulatory genes. Binding to SCO2497, SCO5164 and SCO3185 was conserved across the *Streptomyces* genomes analysed.

Importantly, 14 of the 25 most conserved interactions were previously uncharacterized, revealing a major gap between evolutionary importance and historical discovery. Besides SCO4336, this includes three ArsR-family regulators (SCO6830, SCO1108, SCO6808) and several TFs governing transport systems (SCO3914, SCO7061, SCO1358). The multiDAP dataset also reveals extended regulons of previously studied TFs. For example, the osmotic stress regulator SCO3129 was previously linked to the SCO3128-3130 operon ^49^. However, multiDAP reveals an expanded regulon that includes SCO6717 for acyl-ACP desaturase and the SCO4976-4979 cluster (Fig. 5c), indicating a conserved role in fatty acid metabolism and membrane homeostasis.

At the family level, MerR regulators showed the highest median conservation, while GntR, IclR and LacI families were among the least conserved (Fig. 5d). Since TF families often contain functionally diverse subfamilies ^50^, we examined the TetR family, one of the most diverse in actinomycetes ^51^. KstR2-like TFs, some of which govern specialized catabolic pathways including cholesterol catabolism, showed significantly higher conservation than other TetR subgroups (Fig. 5e). Examples include repression of the phenylacetic acid degradation (*paa*) operon by PaaR (SCO3833) and the leucine/isovalerate utilization (*liu*) pathway by LiuR (SCO2775). These conserved regulons indicate that actinomycete regulatory evolution preserves TF-target pairings for core metabolic functions, while allowing reassignment of targets in more specialized pathways.

### Conserved control of developmental genes

The data reveal novel *cis-trans* relationships between TFs and well-studied genes involved in the control of cell division and development. We thereby used as threshold that a TF-target interaction in *S. coelicolor* should be validated by binding of the same TF to the orthologous genes in at least three additional actinomycetes. For an interaction network, see Fig. S3. The multiDAP experiments verified many known interactions, such as control of *wblE* by Rex ^52^, of *ramC* by RamR ^26^ and of *whiB* by BldO ^53^. Important new regulatory features were identified among others for the *dcw* cluster, which harbours key cell wall and division genes. Besides control by SCO1383, the multiDAP data show that *ftsZ* is likely also controlled by the TetR-family regulator SCO4099. ArsR-family regulator SCO4445 bound to the *ssgB* promoter in all streptomycetes tested; SsgB is a key developmental control protein that recruits FtsZ to the septum sites during sporulation-specific cell division ^54^. Furthermore, MerR-family regulators SCO3694 and SCO7585 bind upstream of *sepG*, which encodes a membrane anchor for SsgB ^55^. This provides important new leads for the control of FtsZ and its interaction partners in actinomycetes. For the control of other genes of interest we refer the reader to StrepTRN.

### TF associations with BGCs are conserved independent of BGC conservation

To determine how regulatory control of BGCs is conserved across actinomycetes, we grouped predicted clusters into gene cluster families (GCFs) using BiG-SCAPE ^56^ and integrated these with TF binding data (Fig. 6a). Analysis of 14 GCFs present in at least four species revealed conserved regulatory interactions in six clusters, involving nine TFs (Fig. 6b). This includes the hopene BGC, controlled by the regulators SCO0376, SCO6770, and the developmental regulator BldC (SCO4091). Notably, the highly pleiotropic BldC also targets the ectoine cluster, linking the timing of BGC expression to specific life-cycle transitions ^57^. Beyond characterized pathways, our results show the pleiotropic regulator AtrA binding within an uncharacterized RiPP-like cluster (BGC 5), providing a target for future elicitation strategies (Fig. 5c).

**Fig. 6:**
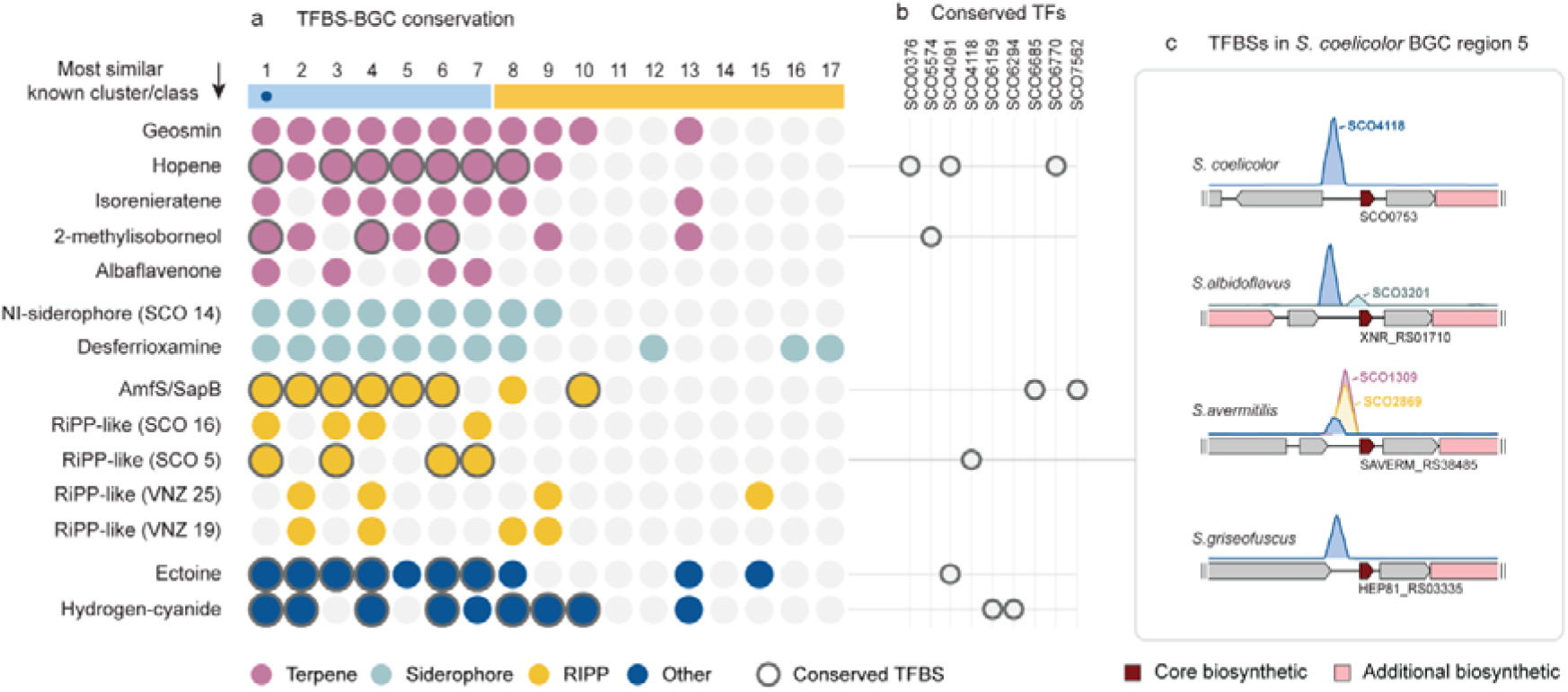
Conservation of regulatory patterns in BGCs across actinomycetes. **a,** Conservation of BGCs among 17 actinomycetes, with *S. coelicolor* indicated by a dark blue dot. The light blue horizontal bar indicates members of the *Streptomyces* clade, while yellow indicates more distantly related species. Colored dots represent conserved BGCs, annotated by BGC type; only BGCs present in at least four species are shown. Clustering is based on BiG-SCAPE. BGCs with evidence of conserved regulation are outlined in grey. **b,** TFs associated with the observed conservation patterns. **c,** TF binding peak distribution for *S. coelicolor* BGC region 5, a RiPP-like cluster.

Next, we examined regulatory mechanisms beyond orthologous clusters, as regulatory associations can persist even when BGC architecture is not conserved. For example, while the bisenarsan BGC is not conserved across the 17 multiDAP species, the four ArsR-like TFs (SCO3696, SCO6808, SCO6812, and SCO6830) bind upstream of a class IV lanthipeptide precursor gene (SGR_RS37785) in *Streptomyces griseus*, suggesting metal-responsive control. Similarly, the zinc-dependent regulator AbsC (SCO5405) targets the core biosynthetic gene of a class III lanthipeptide cluster in *Streptosporangium roseum*. These observations reveal conservation - at least in some cases – of BGC regulation at the level of regulatory inputs rather than gene cluster identity.

## DISCUSSION

The release of the *S. coelicolor* genome sequence some 25 years ago transformed the field by allowing researchers to browse the genome, identify genes and clusters of interest, and formulate hypotheses that were not possible from phenotype alone. Here, we provide an analogous resource for transcriptional regulation. By mapping genome-wide DNA binding for hundreds of transcription factors, we convert the regulatory landscape of *S. coelicolor* from a collection of individually studied regulators into a browsable atlas of transcriptional control. This allows researchers to query, for any gene, operon or pathway, which regulators may control it, and conversely, for any regulator, which genes and processes it may govern. The value of this resource extends beyond specialized metabolism, providing entry points into the regulatory control of development, stress responses, primary metabolism, secondary metabolism, transport, cell division, signaling and the many uncharacterized genes that still dominate the *Streptomyces* genome. Because BGC expression is often embedded in developmental, metabolic and stress-response networks, the atlas also provides a practical route to identify regulatory entry points for activating silent or weakly expressed clusters. Through StrepTRN, these data can be explored directly by gene, regulator and pathway, enabling immediate use by the wider community.

Using DAP-seq across 789 predicted TFs from *S. coelicolor*, we obtained high-confidence binding sites for 393 regulators, increasing the proportion of experimentally supported *S. coelicolor* TFs from ∼8% to ∼50%. This major expansion reveals extensive cross-regulation between core cellular processes and BGCs and exposes network architecture that only becomes visible at this scale. Rather than isolated regulons, the *S. coelicolor* network contains highly connected pleiotropic regulators, many locally acting TFs and dense TF-TF connectivity, together forming a hierarchical and interconnected regulatory system. Benchmarking against previously characterized targets and our *in vivo* ChIP-seq datasets showed strong concordance with published EMSA and ChIP-on-chip data. As anticipated, DAP-seq and ChIP-seq capture complementary aspects of TF binding: DAP-seq enables scalable, condition-independent detection of binding potential, whereas *in vivo* binding reflects physiological context, DNA topology, cofactors and regulatory state.

The atlas also provides training and benchmarking data for computational prediction of regulatory networks. By expanding experimentally supported actinobacterial TF binding profiles, it can strengthen approaches that use TF motifs, genomic context and functional information to predict regulatory interactions, including TF–BGC links, as illustrated by approaches such a COMMBAT ^58^. More broadly, integration of binding data with functional enrichment, orthology and expression profiling provides a route to assign functions to poorly characterized regulators. In this way, genome-scale regulatory data can move anonymous TF annotations towards testable biological hypotheses, while genes and operons of interest can be linked to candidate upstream regulators for targeted validation. It is important to note that DAP-seq and multiDAP measure DNA binding *in vitro* under standardized conditions and therefore cannot capture regulatory states that depend on specific cofactors, ligands, post-translational modifications, DNA topology or chromosomal context. This may explain why several Bld and Whi regulators did not yield significant profiles.

Extending the analysis across 16 additional actinomycetes using multiDAP places regulatory interactions in an evolutionary context. Conserved regulators such as HspR and FtsR, together with many previously uncharacterized TFs, retain core targets across the phylum, whereas other systems show clear lineage-specific changes in TF-target connectivity. This indicates that evolutionary conservation can highlight regulatory relationships that would be difficult to establish on the basis of a single species alone or by phenotype-driven genetics. Beyond evolutionary insight, multiDAP also provides a practical filter for follow-up: recurrent binding of the same TF upstream of orthologous genes across multiple actinomycetes provides comparative support for biological relevance, whereas species-specific interactions may reflect lineage-specific rewiring. It also reveals differences between TF families: some, such as MerR, show high median conservation of binding at orthologous loci, whereas others, including GntR, IclR and LacI, display more plastic regulons. At the level of subfamilies, the pronounced conservation of KstR2-like regulators governing core steps in catabolic pathways illustrates how specific TF subgroups can remain tightly linked to particular metabolic modules across actinomycete evolution.

The dcw cluster and associated division genes illustrate how the atlas can move from global network structure to specific biological hypotheses. MultiDAP links conserved and lineage-specific regulators to genes that encode key components of the division and sporulation machinery, including *ftsZ* and *ssgB*. These connections suggest that divisome assembly and sporulation-specific cell division are embedded in broader transcriptional programmes than previously recognized. They also provide candidate entry points for dissecting how cell division is coordinated with morphological development in streptomycetes.

The same logic applies to specialized metabolism. BGCs are more plastic than many core operons, yet regulatory control can persist even when cluster architecture changes. In such cases, TFs may respond to shared environmental cues while controlling distinct downstream biosynthetic outputs, preserving conserved sensing modules but allowing reassignment of target genes. A recent DAP-seq study that focused on putative gamma-butyrolactone-responsive cluster-situated regulators in *Streptomyces* showed that CSRs may regulate multiple BGCs as well as autoregulatory genes ^23^. Thus, BGC expression is embedded within broader regulatory networks rather than controlled only by local cluster-specific regulators. Conserved regulatory signals may help identify and activate BGCs across diverse genomes, even when the clusters themselves are not conserved. This principle is exemplified by ArsR-family regulators associated with different BGCs across species and by the bisenarsan cluster in *S. coelicolor*, which receives input from several ArsR-family regulators and from the SOS regulator LexA. These links connect metal stress, transport or resistance functions and DNA damage responses to specialized metabolite production. Together, our data argue for a layered view of BGC regulation in which local cluster switches are embedded in broader regulatory hierarchies.

In conclusion, we reconstruct the regulatory architecture of *S. coelicolor* at genome scale and extend this analysis across actinomycetes. By expanding the experimentally supported regulome from ∼8% to ∼50%, we convert a fragmented view of individual regulators into a browsable atlas of transcriptional control. This resource enables researchers to identify candidate regulators for genes, operons and pathways of interest, assign functions to poorly characterized TFs, and explore regulatory conservation and rewiring across species. Beyond serving as a reference map, the atlas provides regulatory entry points for activating silent BGCs and linking specialized metabolism to development, metabolism and stress responses. As genome sequencing revealed hidden metabolic potential and transformed bacterial genetics, large-scale reconstruction of regulatory networks now begins to uncover the control logic that governs these processes, establishing regulatory decoding as a route to understand, predict and engineer complex bacterial phenotypes.

## ONLINE METHODS

### Strains, media, and growth conditions

Bacterial strains are listed in Tables S7 and S8. *Escherichia coli* strain DH5α was used for routine cloning and plasmid propagation. *E. coli* ET12567(pUZ8002) was used for intergeneric transfer of plasmids from *E. coli* to *S. coelicolor* M145 ^59^. *E. coli* strains were grown in Luria-Bertani (LB) medium at 37°C. *S. coelicolor* M145 was the parental strain for all mutants. *S. coelicolor* strains were grown on soya flour mannitol (SFM) for conjugation and sporulation ^59^. Media were supplemented with the appropriate antibiotics when required (apramycin, 50 µg/mL; kanamycin, 50 µg/mL; chloramphenicol, 25 µg/mL; nalidixic acid, 10 µg/mL). For ChIP-seq experiments, approximately 10^7^ spores were cultivated at 30°C in 25 mL liquid minimal medium (NMMP) without PEG ^59^, supplemented with 0.5% (w/v) mannitol and 1% (w/v).

### DAP-seq experiments

DAP-seq experiments were conducted using HaloTagged TF constructs as described in the original DAP-seq protocol ^22^, but using the multiDAP protocol ^21^, with minor modifications. Genomic DNA was sheared to an average size of 75 bp using a Covaris LE220-Plus ultrasonicator and converted into sequencing libraries with the KAPA HyperPrep kit. Before use in DAP-seq, libraries were PCR-amplified for eight cycles to remove DNA modifications. TF-coding sequences were codon-refactored for expression in *E. coli* using BOOST ^60^, applying the *E. coli* codon frequency table to eliminate synthesis constraints (Table S9) and synthesized commercially.

To enable scarless, in-frame fusions with a HALO-tag, 30-nucleotide DNA linkers were appended to both the 5′ and 3′ ends of each sequence. The modified sequences were cloned in-frame downstream of the HaloTag in pIX-Halo_PaqCI, and final constructs were verified by DNA sequencing. Linear HaloTag–TF expression templates were PCR-amplified using the primers pIX-Halo-T7-fwd (5’-GTGAATTGTAATACGACTCACTATAGGG) and pIX-Halo-AfterPolyA-rev (5’-CAAGGGGTTATGCTAGTTATTGCTC) and purified. At least 500 ng of each linear PCR product was used for *in vitro* protein synthesis using the TnT T7 Quick for PCR DNA kit (Promega). Each DNA affinity purification reaction was run with 50 µL expressed protein, 1 ng of the previously prepared genomic DNA fragment libraries, and 1000 ng salmon sperm DNA as a non-specific competitor to reduce non-specific binding. DAP reactions were performed with 50 µl of expressed HaloTag–TF protein, 1 ng of genomic DNA library and 1 µg salmon sperm DNA as nonspecific competitor. The final DAP-seq libraries were sequenced with a NovaSeq 6000 or NovaSeq X Plus (Illumina), targeting 4 million 2×150 reads per sample.

### Peak data processing pipeline

Sequenced reads were adapter trimmed and quality filtered using BBTools v38.90 ^61^ using the following options: k=21 mink=11 ktrim=r tbo tpe qtrim=r trimq=6 maq=10. Filtered reads were aligned to the reference genome using bowtie2 v2.4.2 ^62^ with the options --no-mixed --no-discordant. Peaks were called using MACS3 v3.0.0a6 ^63^, using combined negative control samples with mock protein expression as the background file, and the following options: --call-summits --keep-dup 1, and --gsize with the total reference genome size. Sequences representing up to the 100 strongest peaks, as scored by signal value in column 7 of the narrowPeak files, were extracted using a custom python script. Peak distribution was obtained using MACS3 pileup files. These peak sequences were used to generate motifs using MEME v5.5.7 ^64^ with a zero-order background file generated from the reference genome, and the following additional options: -evt 0.05 -nmotifs 3 -minw 5 -maxw 30 and visualized using Logomaker v0.8 ^65^. Motif calling was run twice for each dataset, once with the entire peak regions and once with only the regions +/- 30 bp flanking the summit location of each peak. Peaks were assigned to putatively regulated genes with BEDTools v2.30.0 ^66^ ^66^, using bedtools subtract to filter for intergenic peaks, followed by bedtools closest to assign each peak to up to two genes immediately adjacent to and downstream of the peak. Peaks were retained only if they showed at least a 30-fold enrichment over background, had their peak start located within 200 bp upstream of a gene start site and had a q-value < 0.001 based on the ‘local Poisson’ model and Benjamini-Hochberg (BH) FDR correction of MACS3.

### ChIP-seq experiments

#### Constructs and cloning

Oligonucleotides and plasmids used in this study are listed in Tables S10 and S11. DNA amplification, cloning and construct verification were performed essentially as described previously, with minor modifications ^67^. Target genes were fused to an N- or C-terminal 3×FLAG tag and expressed from their native promoter. The position of the tag was chosen based on the predicted location of the helix–turn–helix domain, with the distal terminus selected whenever possible to minimize interference with DNA binding.

For N-terminal fusions, the native promoter region and coding sequence were amplified separately, allowing insertion of the ATG–3×FLAG sequence immediately upstream of the coding region, and cloned into BP001. For C-terminal fusions, the promoter region and coding sequence lacking the stop codon were cloned into BP001 or pCOMM001, either together with a separately amplified 3×FLAG fragment or directly into pCOMM001, which contains the 3×FLAG sequence downstream of the cloning site. Correct assembly of all constructs was confirmed by Sanger sequencing.

#### DNA-protein cross-linking and chromatin immunoprecipitation

ChIP-seq was performed essentially as described previously for FLAG-tagged proteins in *S. coelicolor* ^67^. Approximately 10□ spores of COMM006 or 10□ spores of COMM021, COMM029, COMM031, COMM034, COMM055, COMM056, COMM057, COMM060, COMM061 and COMM066 were inoculated in 25 ml NMMP medium and grown for 46 h. Cultures were cross-linked with 1% formaldehyde for 10 min at 30°C, followed by quenching with glycine. Biomass was collected, washed with ice-cold TBS and lysed enzymatically before chromosomal DNA was sheared to fragments of approximately 100–500 bp using a Bioruptor Plus sonication system.

FLAG-tagged protein–DNA complexes were immunoprecipitated overnight at 4°C using anti-FLAG M2 magnetic beads. A fraction of the cleared lysate was retained as input genomic DNA control. After washing, protein–DNA complexes were eluted, cross-links were reversed and DNA was purified using the DNA Clean & Concentrator-5 kit. ChIP and input DNA samples were sequenced by BGI using the DNBSEQ-G400 platform in single-end 50-bp mode.

### Electrophoretic Mobility Shift Assays

Oligonucleotide sequences used for EMSA are described in Table S9. *S. coelicolor* genomic DNA was used as the template to PCR-amplify DNA fragment containing the SCO1383 open reading frame. The fragment was subsequently cloned into vector pET21a (Novogen) to express SCO1383 with a C-terminal 6×His tag. Protein purification was performed according to the supplier. As probe we used the −140/−81 region relative to the *ftsZ* translational start site, whereby oligonucleotides PZ_EMSA_F and PZ_EMSA_R were used to amplify the DNA fragment from the *S. coelicolor* genome. As a negative control, a 60 bp random sequence probe was generated by annealing oligonucleotides Random_EMSA_F and Random_EMSA_R. EMSAs were performed as described ^30^.

### Reference genomes

The reference genome sequences of the strains were retrieved from the NCBI Reference Sequence Database (RefSeq), with accession numbers provided in Table S7. Phylogenetic analysis was performed using PhyloPhlAn v3.0.60 ^68^ with DIAMOND v2.0.6 as a mapping tool, MAFFT v7.475 for the multiple sequence alignment, trimAl v1.4.rev15 for alignment trimming and IQ-TREE v2.0.3 with 100 bootstrap replicates for phylogenetic tree building. The protein domain architecture of each TF was determined using the regulatory PFAM subset library and the approach described in previous work ^50^.

### Functional assignment of TFs

Functional gene annotations were assigned using the EggNOG-mapper v2.1.12 ^69^. Enrichment was assessed via Mann-Whitney U tests with BH FDR correction using a q-value threshold of□0.05. The results were visualized using Matplotlib ^70^. For pairing the TF binding sites with expression data, we obtained the RNA-seq data reported by ^37^ and used the transcripts per million and log_2_-transformed expression from xylose- and glucose grown cultures. Statistical analysis of differential gene expression between the two conditions was performed using GraphPad, which implements an unpaired t-test and FDR correction using the Benjamini-Krieger-Yekutieli approach.

### BGC analysis

For each reference genome, BGCs were detected using the prediction software antiSMASH v7.1.0 ^38^ with the detection strictness set to “relaxed.” BGC boundaries were defined based on experimentally validated ranges from the MIBiG v3.0 ^71^ database when the BGC had a MIBiG similarity score of at least 90%. In cases where no match to a validated BGC was found, the operon structure containing the antiSMASH-assigned biosynthetic core genes was used, allowing a maximum intergenic distance of 200 bp. The clustering algorithm BiG-SCAPE v1.1.8 ^56^ was used to group the BGCs into groups of similar gene clusters (*i.e.* gene cluster families, GCFs). For BGCs that only comprise a single or few biosynthetic gene(s), their flanking regions can dominate the span of the antiSMASH-detected regions and therefore cause these BGCs to appear as singletons rather than being grouped into their respective GCFs. Therefore, the resulting clustering was manually inspected, and any misclassified BGCs were reclassified into their correct families based on their MIBiG or KnownClusterBlast similarity score. TF-BGC associations were reported when the peak summit fell within the regulatory region, defined as -350 bp to +50 bp relative to the start codon of each gene.

### Conservation study

To assign individual genes to orthologous groups from 17 actinomycetes, we used OrthoFinder2 v2.5.5 ^72^ with default parameters. If multiple genes from the same species are assigned to a single orthogroup, each gene was weighted based on the number of genes from that species within the orthogroup. To analyze conservation of target genes, we generated 10,000 sets of randomly selected genes of equal size per species for each target gene set and calculated match scores by comparing the overlap between observed and random sets. P values were determined by calculating the fraction of random comparisons with match scores equal to or greater than the observed match score. Target gene conservation between species was assessed by counting the number of species in which a given TF also targeted at least one gene within the same orthogroup. This count was subsequently converted into a percentage relative to the total number of species considered.

## ACKNOWLEDGEMENTS

The project was supported by Advanced Grant 101055020-COMMUNITY from the European Research Council to G.P.v.W., Starting Grant 948770-DECIPHER from the European Research Council to M.H.M., and by the U.S. Department of Energy Joint Genome Institute (proposal 10.46936/10.25585/60008131; https://ror.org/04xm1d337), a DOE Office of Science User Facility, supported by the Office of Science of the U.S. The Department of Energy operated under Contract No. DE-AC02-05CH11231. HA was supported by the donors of the Leiden University Fund, Swaantje Mondt Fonds.

