## Supplemental Figures for "A global regulatory atlas of *Streptomyces* reveals conserved and diversified transcriptional networks across actinomycetes"

Belonging to the manuscript

* These authors contributed equally

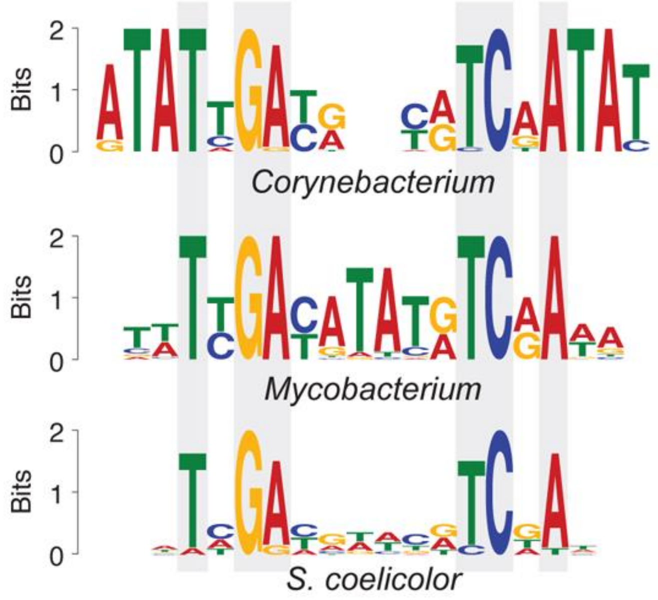


**Figure S1.** Comparison of ArsR binding motifs from *Corynebacterium* and *Mycobacterium* species of RegPrecise [1], with the DAP-seq SCO3696 *Streptomyces coelicolor* motif.

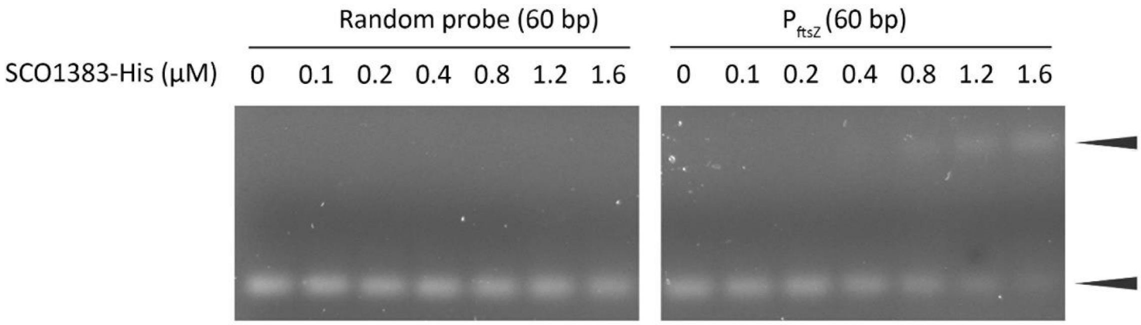


**Figure S2. SCO1383 binds to the *ftsZ* promoter region.** EMSAs showing interactions between SCO1383 and a partial *ftsZ* promoter sequence (−140 to −81 relative to the *ftsZ* translational start site). SCO1383 was incubated with approximately 80 ng of DNA probes, with protein concentrations as indicated. Bottom fragments, unbound DNA; top fragments (lower mobility), protein–DNA complexes.

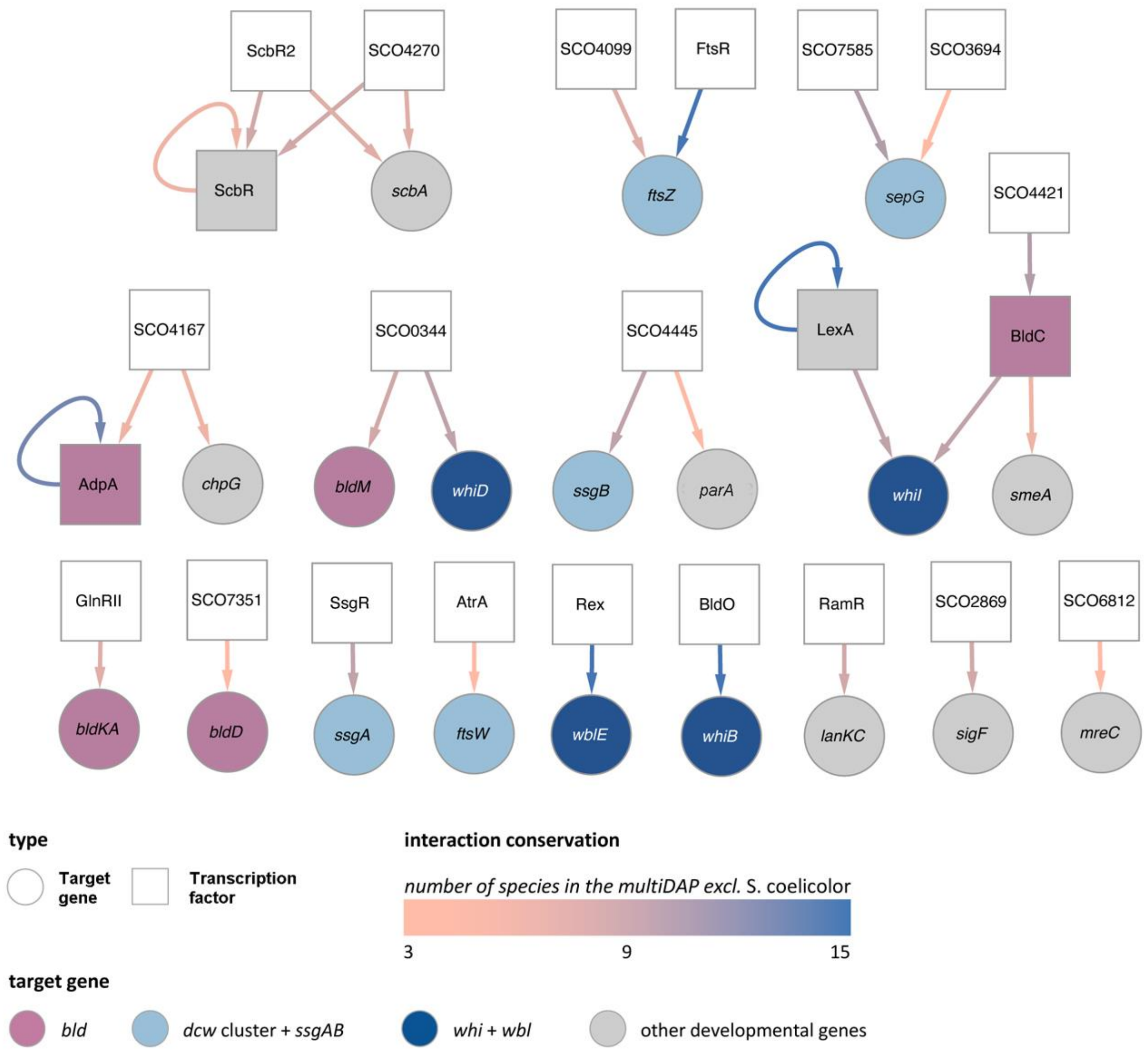


**Figure S3. Conserved regulatory network of developmental genes of *S. coelicolor*.** Interactions between transcription factors (squares) and their target genes (circles) in *S. coelicolor* are shown in the network when multiDAP showed binding of the *S. coelicolor* TF to at least three orthologues in other actinobacteria. Note that target genes can be transcription factors themselves. Auto-regulatory interactions indicate the protein bound in front of its gene. Target genes are colored based on the developmental gene category, while interactions are colored based on the number of species with conserved interaction in the multiDAP. The network was visualized using *Cytoscape v3.10.3* [2].
